# YAP signaling promotes resistance to MEK and AKT inhibition in *NF1*-related MPNSTs

**DOI:** 10.1101/2025.06.16.659334

**Authors:** Lauren E. McGee, Jamie L. Grit, Curt J. Essenburg, Sophia Agrusa, Elizabeth A. Tovar, Lisa Turner, Katelyn Becker, Jason Zbieg, Anwesha Dey, Jennifer E. Klomp, Jeffrey A. Klomp, Marie Adams, Ian Beddows, Emily Wolfrom, Zhen Fu, Angela C. Hirbe, Carrie R. Graveel, Matthew R. Steensma

## Abstract

Neurofibromatosis type 1 (NF1) is a tumor predisposition syndrome caused by loss of function of the neurofibromin protein. Malignant peripheral nerve sheath tumors (MPNSTs) are a rare and deadly sarcoma with few therapeutic options that are the leading cause of death for patients with NF1. To date, no targeted therapies have been approved for MPNST treatment, highlighting the need for an understanding of adaptive signaling mechanisms that drive resistance. We developed a preclinical model of drug resistance using a cross-over drug holiday design and evaluated patterns of response and resistance to MEK and AKT inhibitors, two pathways that are dysregulated by loss of neurofibromin. We show that the mTOR and YAP/TEAD pathways are activated by MEK inhibitor exposure, yet blockade of these pathways in resistant MPNST PDX models does not significantly reduce tumor growth, despite strong *in vitro* synergy between trametinib and the novel TEAD inhibitor, GNE-7883. Using spatial transcriptomics, we uncovered phenotypic inertia as a key mechanism of drug resistance in MPNST, in addition to signaling plasticity. Further, we found that resistance is mediated by sustained ERK, YAP, and MYC driven transcriptional programs. In the future, preclinical studies should focus on addressing intratumoral heterogeneity and how it evolves over time.

## Introduction

Neurofibromatosis Type 1-related malignant peripheral nerve sheath tumors (NF-related MPNSTs) are aggressive sarcomas that are highly resistant to chemotherapy (1-3). MPNSTs are common in NF1, occurring in 8-13% of individuals (4). NF1 is characterized by germline NF1 haploinsufficiency which results in diminished or absent levels of intracellular neurofibromin. Because neurofibromin is a critical upstream regulator of RAS signaling, both MPNSTs and their haploinsufficient stroma are subject to deregulated RAS signaling. Despite an apparent dependency on RAS/MAPK signaling, efforts to target the RAS/MAPK pathway in NF-related MPNSTs have been associated with profound therapy resistance due, in part, to kinome adaptation (5). MEK inhibitors, in particular, have shown promise in preclinical models of NF-related MPNSTs, however clinical trials failed to show response to single agent or combination therapies. Because MPNSTs frequently exhibit a large burden of genomic alterations and clonal heterogeneity, they are prone to rapid development of resistance (6-8). There are currently no targeted therapies approved for NF-related MPNSTs, highlighting the need for a better understanding of the complex signaling in these tumors and how mechanisms of resistance arise (2, 9).

The single agent MEK inhibitor, selumetinib, is FDA approved for the treatment of benign, plexiform neurofibromas (PNFs), but has failed to show long term success in cancers dependent on hyperactivated RAS signaling including MPNST, colon cancer, and melanoma (10). We recently showed that AKT activation was a unifying mechanism of MEKi resistance among three genomically diverse, *NF1*-deficient MPNST GEMM models (6, 11). Further studies demonstrated compensatory kinase signaling activation after inhibition with trametinib, including increased PDGFRB, AKT, and CRAF-related pathways (12).

More recently, studies have implicated Hippo signaling in MPNST progression and therapy resistance. The Hippo pathway is a key regulator of organ growth and cellular differentiation. Hippo functions as a tumor suppressor through phosphorylation of the effector protein YAP1, which leads to YAP1 sequestration in the cytoplasm (13). Upon deactivation of members of the Hippo pathway through external factors such as cell-density, matrix stiffness, mutation, or interaction with other pathways, YAP1 is de-phosphorylated and enters the nucleus to bind with the transcription factor TEAD. YAP1-TEAD binding activates transcription of known target genes such as *MYC*, *CTGF*, and *Cyr61* raising the question whether MPNST therapy resistance is TEAD dependent (14-17). Genomic alterations of YAP and other Hippo family members are rare in NF-MPNSTs; however, Hippo is known to interact with several other pathways, including RAS and AKT, that are capable of altering expression of Hippo family members, including YAP1 itself. YAP1 has been shown to directly interact with RAS and AKT signaling (18-23). Hyper-activated YAP1, through deletion of LATS 1/2 in Schwann cells, was shown to be sufficient to cause MPNSTs in mice (17). In addition, in a panel of soft tissue sarcomas stained for YAP expression, MPNST tissues had the largest proportion of nuclear-localized YAP expression (16). In other cancers, including neuroblastoma and pancreatic cancers, YAP1 upregulation after inhibition of the RAS pathway contributed to therapeutic resistance and metastasis (18-24). Another study utilizing a CRISPR/Cas9 screen to identify vulnerabilities in MPNSTs implicated upregulation of the Hippo pathway as having a role in initiation of neurofibromas (25). Thus, there is sufficient rationale to target YAP-TEAD in NF-related MPNSTs.

Hippo pathway inhibitors are being investigated in a variety of cancer contexts (14, 26). The novel pan-TEAD inhibitor, GNE-7883, prevents the interaction between YAP/TAZ and all four TEAD paralogs to prevent chromatin accessibility at TEAD target genes. GNE-7883 has been shown to decrease proliferation in cancer cells with altered YAP/TAZ signaling (27). Furthermore, GNE-7883 successfully reduced tumor volume and viability in cancers with resistance to the KRAS G12C inhibitor, sotorasib, proving its ability to target YAP/TAZ activation after RAS pathway inhibition (27). Here, we not only investigate the role of AKT inhibitors in MEK inhibitor resistance, but we also use GNE-7883 to target downstream YAP activation in MEKi-resistant, NF-related MPNSTs. Our data demonstrates synergy between TEAD and MEK inhibitors across resistant and parental MPNST cell lines, as well as new mechanisms of MEK inhibitor resistance in MPNST patient-derived xenograft (PDX) models, highlighting the critical role of YAP in modulating RAS and AKT signaling.

## Results

### Combined MEK and AKT inhibition is ineffective in MPNST cells

Previously, we demonstrated that AKT activation led to MEK inhibitor resistance in genetically diverse mouse models of *NF1*-related MPNSTs (11). To determine if this mechanism of resistance was present in human MPNST cell lines, we treated NF90 and S462 human MPNST cell lines with the MEK inhibitor trametinib for 2 and 24 hrs. Western blot analysis of pAKT S473 and pERK T202/Y204 confirmed that this adaptive response to MEK inhibition also occurs in the human MPSNT cell lines within two hours, and up to 24 hours after treatment (Figure 1A). To determine if combined AKT and MEK inhibition would have additive or synergistic inhibition in human MPNST cells, we treated cells with trametinib and the pan-AKT inhibitor, afuresertib, in a matrix dose response assay. Combined trametinib and afuresertib resulted in no additive or synergistic effect on cell viability (Figure 1B and C, Supplemental Figure 1A), although single agent MEK inhibition was modestly effective. On target effects were confirmed by western blot, with loss of pERK in trametinib treated cells, and increased pAKT in afuresertib treated NF90 cells (Figure 1D), likely due to loss of negative feedback (28).

**Figure 1:**
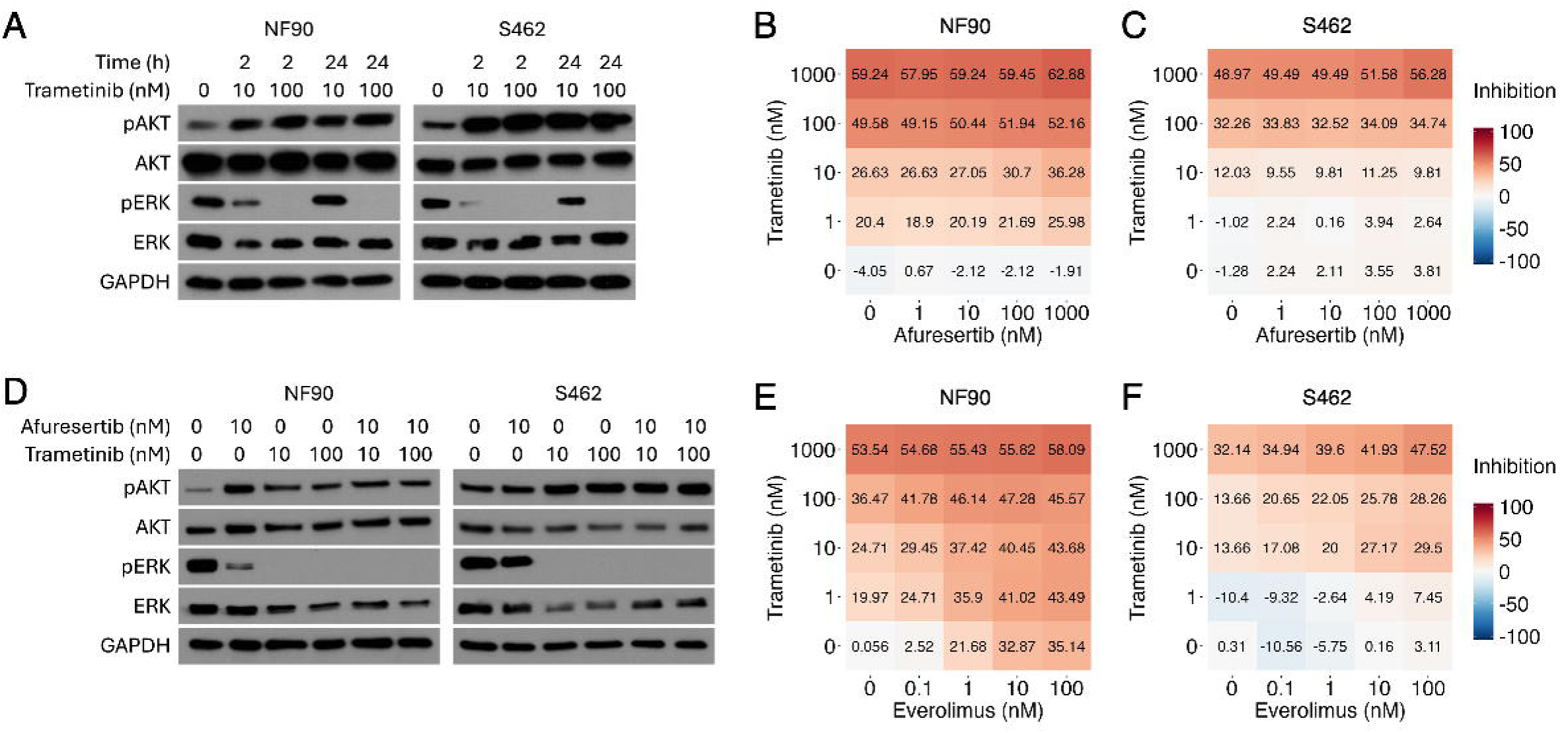
Combined MEK and AKT inhibition does not abrogate MPNST cell growth A) Western blot analysis of AKT and ERK signaling with trametinib treatment in NF90 and S462 cells treated with 100 and 10 nM trametinib for 24 or 2 hours. B) Dose response matrices of NF90 and C) S462 percent inhibition in response to combined treatment with trametinib and afuresertib. D) Western blot analysis of AKT and ERK signaling after treatment with 10 and 100 nM trametinib, and/or 10 nM afuresertib. E) Dose response matrices of NF90 and F) S462 percent inhibition in response to combined treatment with trametinib and afuresertib.

Given the lack of cell death to afuresertib, we sought to determine whether MPNST cell lines respond to the pan-AKT inhibitor, ipatasertib, with or without trametinib. Ipatasertib had no effect on viability as a single agent, (Supplemental Figure 1B) and no synergy was present between ipatasertib and trametinib (Supplemental Figure 1C). Next, we treated the MPNST cell lines with the mTOR inhibitor, everolimus, to determine if mTOR dependency was present in the absence of AKT inhibition. Dose-dependent treatment responses were observed for both trametinib and everolimus in the NF90 cell line, but S462 cells were resistant to single agent everolimus (Figure 1E and F). Interestingly, the combination of everolimus and trametinib was synergistic in the S462 line (max Bliss synergy score of 13.37) (Supplemental Figure 1D). Together, these results show that inhibition of AKT alone or in addition to MEK inhibition is ineffective in reducing MPNST viability, whereas downstream mTOR inhibition cooperates with MEK inhibition *in vitro*.

### Altered signaling patterns with MEK and mTOR inhibition in a drug holiday treatment model

As shown by our lab and others, the molecular and genetic heterogeneity of MPNSTs leads to rapid kinome adaptation during treatment (6, 8, 12, 25, 29). Short term treatment responses in MPNST are commonly demonstrated in cell line-based models and GEMMs. Human PDX models are more representative of human disease, having preserved the original clonality of the source tumors (8). To determine adaptive and acquired resistance mechanisms in response to trametinib and/or everolimus therapy, we utilized highly resistant MPSNT PDXs. That is to say that we attempted to select the PDX models with the most resistant clonal populations to study how kinome adaptation evolves over time in a solid tumor.

To identify alternate pathways influencing MPNST resistance, we induced kinome adaptation by modulating the timing of drug delivery. We employed a unique experimental design to determine patterns of kinase signaling in response to both primary and secondary targeted therapies (Figure 2A). This cross-over, “drug holiday” dosing schedule allowed for combined assessment of tumor kinetics and intra-tumoral kinase signaling over time. When MPNST PDX tumors reached approximately 150 mm^3^ (T_0_), they were treated for 15 days with everolimus (mTOR), trametinib (MEK), or combined everolimus + trametinib (mTOR + MEK). After 15 days of treatment, animals started a drug holiday period (T_1_) until the tumor growth increased >1000 mm^3^ (T_2_). At this point, mice resumed therapy with either everolimus and/or trametinib until tumors reached euthanasia criteria at 2500 mm^3^ or had been on treatment for 15 days (T_3_) (Figure 2A). This strategy allowed us to not only measure initial treatment response but also measure the tumor response to drug removal and subsequent perturbation with a secondary treatment. In addition, we assessed response to re-exposure of the same inhibitor (i.e. trametinib ➔ trametinib) after a drug holiday vs. switching to a different pathway inhibitor (trametinib → everolimus). We utilized the MPNST PDX MSTA-440-2 for these studies, which represents a heavily pretreated MPNST that arose from an NF1 subject being treated with selumetinib for a large plexiform neurofibroma.

**Figure 2:**
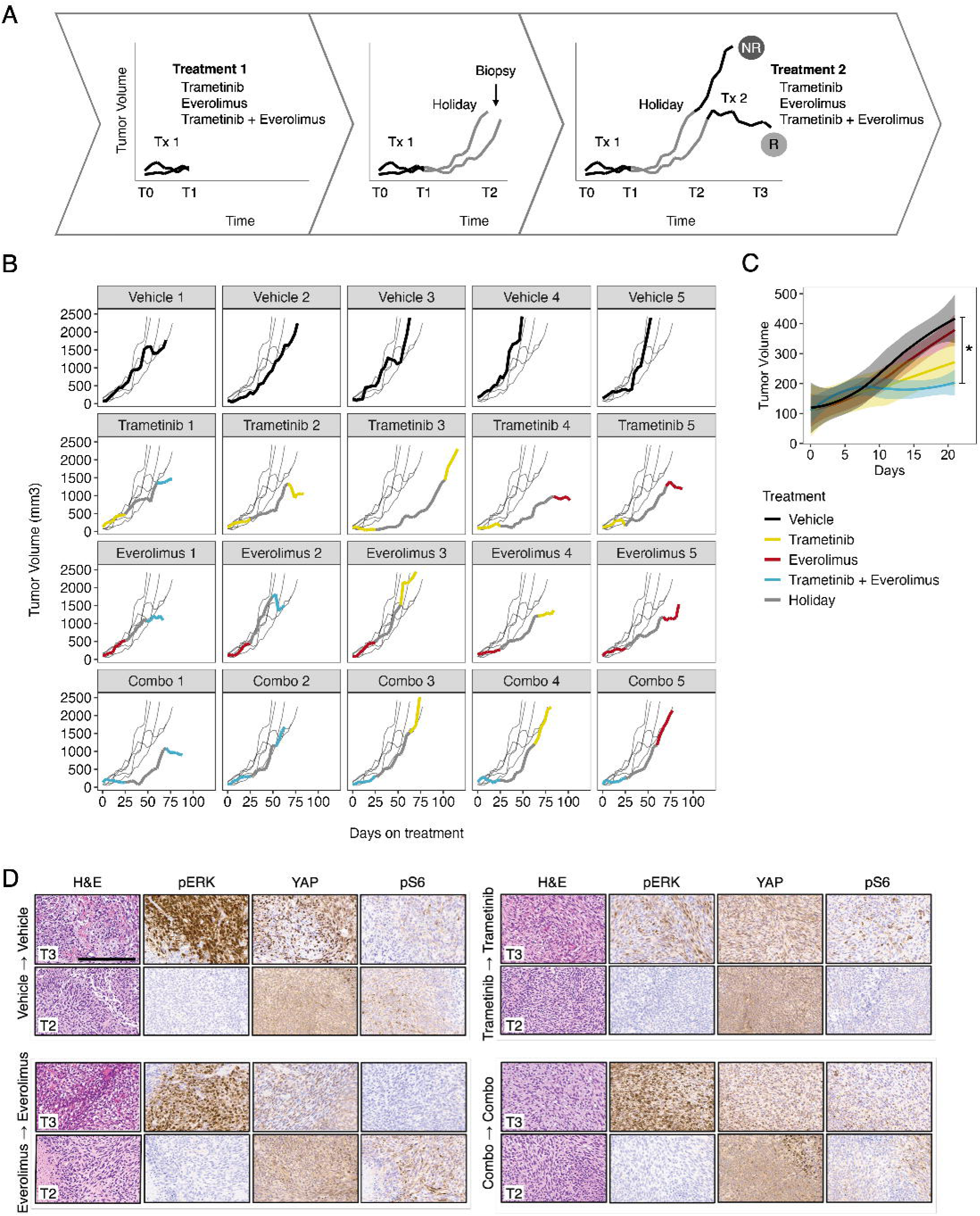
PDX model of MPNST reveals heterogenous response to MEK and mTOR inhibition A) Schematic of cross-over and drug holiday treatment model. Tumorgrafts were monitored until they reached 150 mm^3^ (T_0_), at which point animals were randomized and placed on Treatment 1 (Tx1; 1 mg/kg trametinib, 5 mg/kg everolimus, or a combination) for 3 weeks. At this timepoint (T_1_), animals were placed on drug holiday until their tumors reached 1000 mm^3^ (T_2_) and then were biopsied and randomized to Treatment 2 (Tx2; either a new treatment or their previous treatment). Tumorgrafts were monitored until non-responders (NR) reached euthanasia criteria or responders (R) received 15 days of treatment (T_3_). B) Individual growth curves of MSTA-440-2 PDX lines colored by treatment and plotted against the vehicle tumor growth curves (black). C) Loess smoothed growth curves for Treatment 1. D) Representative IHC of pERK, YAP, and pS6 of re-exposure treated MSTA-440-2 PDXs lines at T_2_ and T_3_. * adjusted p value < 0.05

First, we evaluated primary treatment response compared to vehicle-treated tumors. As expected, neither single agent everolimus or trametinib significantly altered tumor kinetics when used as primary treatment (Figure 2B & C). The initial treatment of the combination of trametinib and everolimus did, however, effect significantly slower growth when compared to vehicle (Figure 2C). Notably, several mice that were initially treated with trametinib (as single agent or in combination with everolimus) demonstrated delayed tumor growth after being placed on holiday. However, following this delayed rebound, rapid tumor growth resumed in only a matter of days (Figure 2B), consistent with an adaptive kinome response.

When we evaluated the response to secondary treatment regimens, we observed heterogeneous patterns of resistance that were partially dictated by pretreatment. For example, 4/5 tumors that were initially treated with combination therapy were profoundly resistant to subsequent treatments (Figure 2B). In contrast, tumors that were initially treated with single agent therapy generally showed good response to secondary therapies, although this was not statistically significant, possibly due to the small sample numbers in each group upon crossover treatment and response heterogeneity. We also evaluated the MEKi/mTORi drug holiday paradigm in the human MPNST PDX line WU-356 with similar results, although several of these tumors grew too quickly throughout primary treatment and drug holiday to use for the extended treatment study (Supplemental Figure 2).

To identify potential cell signaling adaptations that drive the heterogeneous responses to targeted inhibition, tumors were immunostained for pERK and the mTOR activation marker S6 at timepoints T_2_ and T_3_. In addition, we examined YAP signaling due to previously reported literature characterizing YAP as a route of resistance to MEK inhibition (18, 20, 21, 30-32). Immunostaining revealed heterogenous expression of pERK, S6, and nuclear YAP in both the MSTA-440-2 and WU-356 tumors, regardless of treatment and timepoint (Figure 2D, Supplemental Figure 3), suggesting that regional MEK/mTOR pathway reactivation likely contributes to therapy resistance. Additionally, in several treated tumors we identified strong upregulation of nuclear YAP in regions with low/absent ERK staining (Supplemental Figure 3). Collectively, these data indicate that distinct populations within PDX tumors leverage alternate signaling pathways to drive tumor growth, with a potential switch occurring between ERK and YAP signaling that may act as a node of resistance in these tumors.

To further investigate the molecular changes occurring in PDX MPNSTs in response to treatment, we utilized the NanoString GeoMx Digital Spatial Profiler (DSP) platform for spatial transcriptomics of pERK and YAP positive tumor regions. We selected proliferative (Ki67 positive) regions of MSTA-440-2 tumors at both the T_2_ and T_3_ timepoints for transcriptional profiling of ERK positive, YAP positive, and double negative segments (Figure 3A & B). Gene set enrichment analysis (GSEA) revealed significant upregulation of Hallmark MYC Targets and Epithelial Mesenchymal Transition (EMT) gene signatures at the T_3_ timepoint, regardless of vehicle or drug treatment (Figure 3C & D, Supplemental Figure 4A, and Supplemental Table 1). This coincided with deregulation of multiple oncogenic signaling pathways, most notably KRAS signaling (Figure 3E, Supplemental Figure 4B, and Supplemental Table 1). Interestingly, even after controlling for nuclear YAP expression, the only oncogenic signaling pathway that was consistently upregulated at the T_3_ timepoint was the YAP Conserved Signature established by Cordenosnsi *et al*. (Figure 3F, Supplemental Figure 4A, and Supplemental Table 1). Remarkably, enrichment of the YAP signature occurred in both YAP positive and negative regions (Figure 3G).

**Figure 3:**
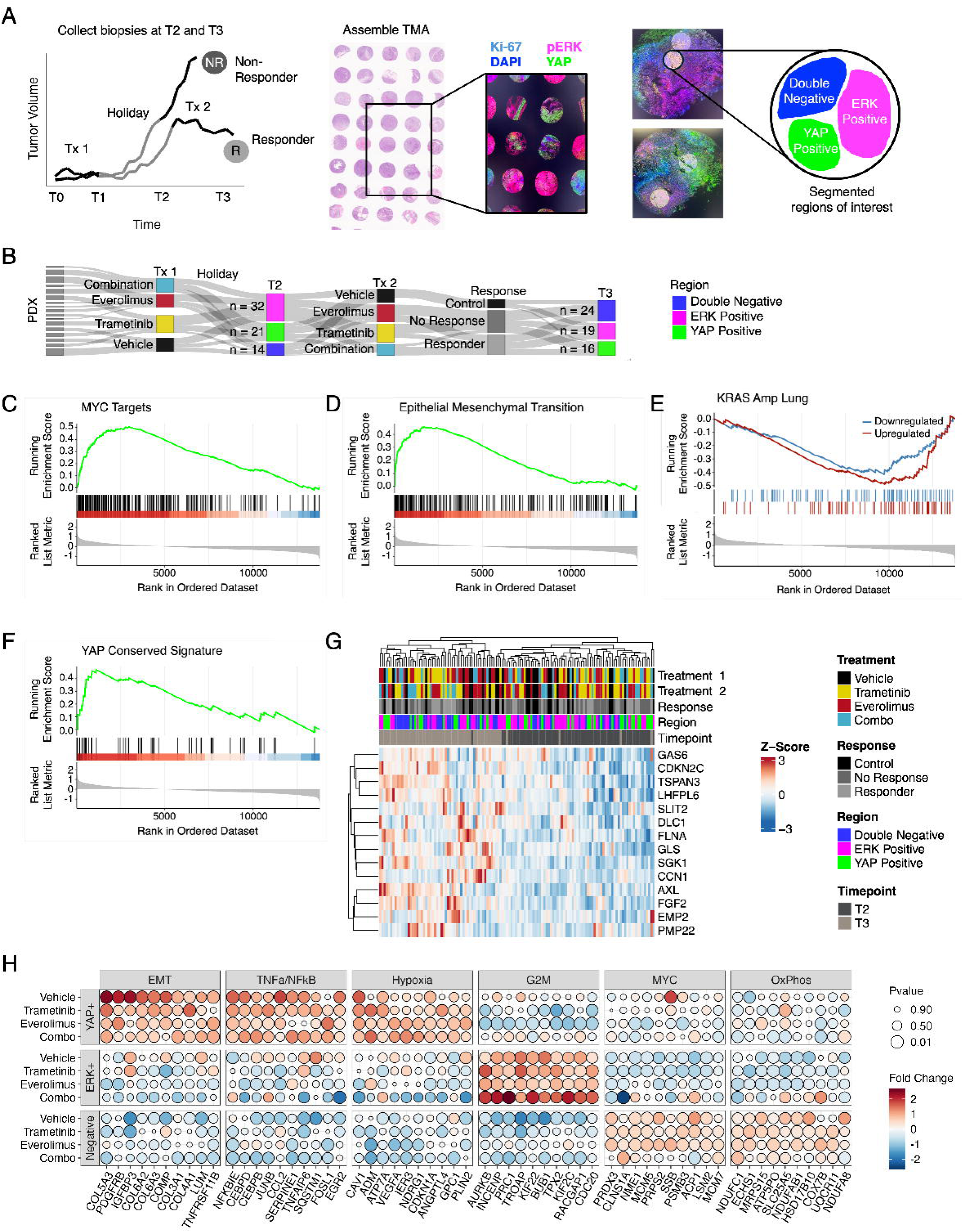
Spatial transcriptomics identifies functionally distinct cell populations in MPNST PDX A) GeoMx experimental approach. T_2_ and T_3_ biopsies from Figure 2 were assembled into a tissue microarray and stained for IF with DAPI and Ki-67 markers to identify proliferative regions of interest, which were further segmented based on pERK and nuclear YAP expression and then sequenced. B) Sankey diagram showing the number of segmented regions sequenced for each timepoint. C) GSEA plots for MSigDB Hallmark MYC Targets and D) EMT gene sets, and E) MSigDB Oncogenic KRAS and F) YAP Signature gene sets at the T_3_ timepoint compared to T_2_. G) Heatmap of the core enrichment genes in the YAP Conserved signature annotated by treatments, response, region, and timepoint. H) Dotplot of the T_3_ top 10 core enrichment genes in each Hallmark pathway, stratified by segmented region and Treatment 2, sized by p value, and colored by log2 fold change in expression.

To understand how YAP signaling contributes to drug response, we evaluated differentially expressed genes (DEGs) between YAP positive and YAP negative tumor regions at the T_3_ timepoint. Interestingly, EMT gene upregulation was YAP dependent, regardless of drug treatment (Figure 3H, Supplemental Table 1). This is consistent with the role of YAP in other tumor types, where it promotes EMT and metastatic progression (18, 19). Inflammatory pathways and hypoxia were also enriched in the YAP positive population (Figure 3H, Supplemental Table 1). We next hypothesized that ERK positive regions contributed to MYC target gene expression, as MYC is downstream of ERK, however, GSEA revealed no upregulation of the MYC Targets signature in ERK positive regions (Figure 3H, Supplemental Table 1). Rather, we identified treatment-specific upregulation of the G2M Checkpoint, Mitotic Spindle, and E2F Targets gene sets in association with deregulation of multiple oncogenic signaling pathways (Figure 3H, Supplemental Table 1). We next examined tumor regions that were negative for both YAP and ERK, and found enrichment of the MYC Targets, suggesting the presence of 3 transcriptionally distinct cell populations (Figure 3H, Supplemental Table 1). Oxidative phosphorylation genes were also enriched in this population (Figure 3H, Supplemental Table 1). Notably, the MYC signature was absent in double negative regions from tumors that received combination trametinib and everolimus at treatment cross-over (Figure 3H, Supplemental Table 1), of which 3/4 were considered responders. Collectively, these data suggest that transcriptional heterogeneity contributes to treatment resistance and aggressive tumor progression in MPNST.

### Trametinib resistant cells have increased YAP signaling

Identifying the transcriptional programs that promote MEK inhibitor resistance is especially critical as more MPNSTs will likely develop in the context of MEK inhibition since the approval of selumetinib for the treatment of the benign precursor plexiform neurofibroma in 2020. To further examine the molecular mechanisms by which YAP and MYC contribute to treatment resistance, we developed a trametinib resistant MPNST cell line by exposing NF90 cells to low dose trametinib for approximately 5 months, generating the trametinib resistant NF90 (NF90-R). NF90-R cells were resistant to trametinib *in vitro* (Figure 4A) and exhibited more invasive growth in Matrigel than the parental NF90 cell line (Figure 4B), suggesting that chronic trametinib exposure may select for a more invasive phenotype in MPNST. A sustained loss of pERK was also seen in NF90-R cells compared to the NF90 parental line while pAKT expression was largely unchanged between cell lines (Figure 4C).

**Figure 4:**
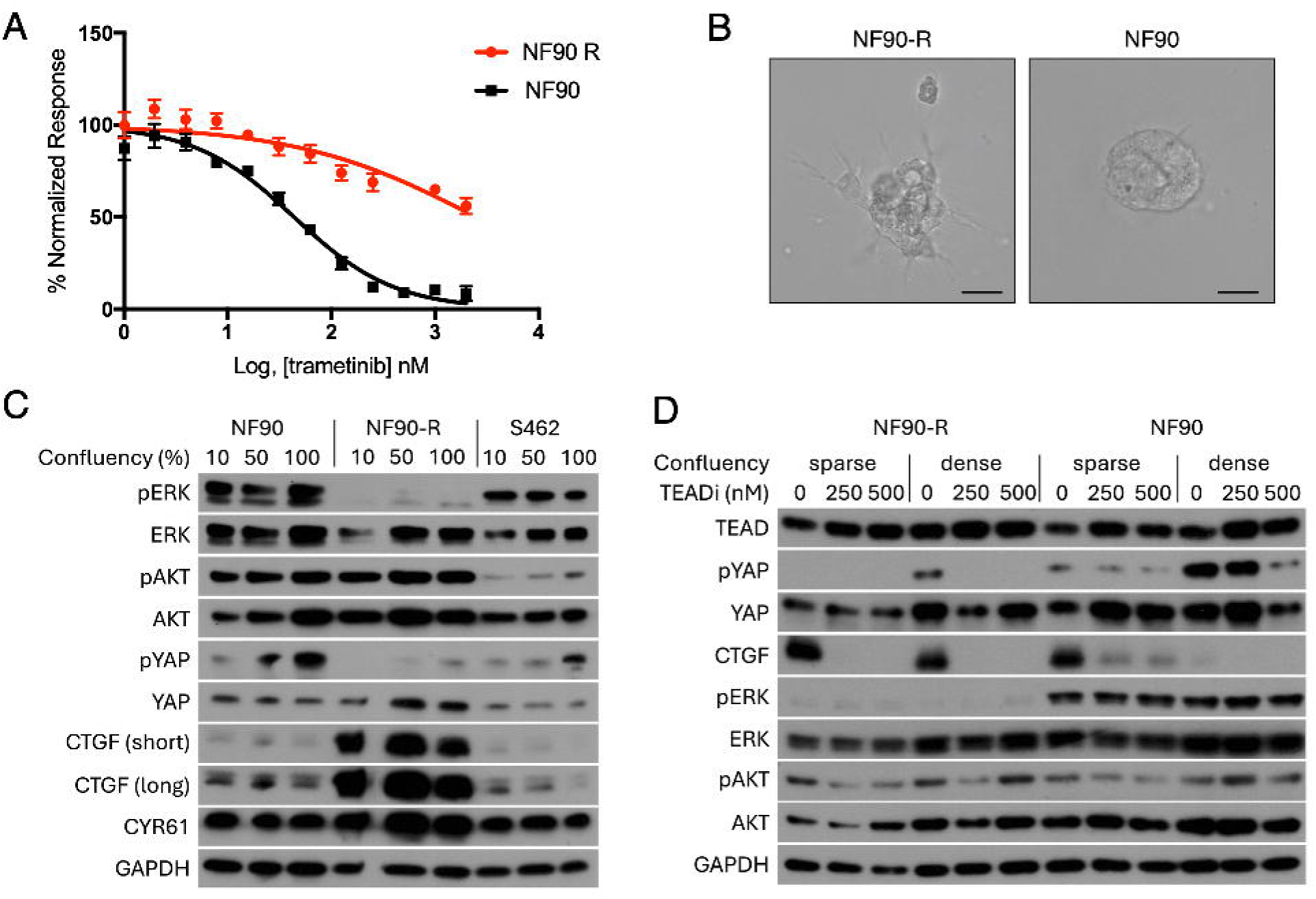
Chronic trametinib exposure promotes cell invasion and YAP/TEAD signaling A) Dose response curve of trametinib in NF90 parental compared to NF90-R cells (NF90 IC50 = 100 nM, the NF90-R IC50 = 800 nM). B) NF90 and NF90-R cells plated on Matrigel for 24 hours, scale bars = 25 um. C) Western blot analysis of NF90, NF90-R, and S462 cell lines characterizing alterations in ERK/AKT and YAP/TEAD signaling at difference plating densities. D) Western blot showing YAP/TEAD and ERK/AKT signaling changes in NF90-R and NF90 cell lines after treatment with TEADi for 48 hours.

We next evaluated YAP pathway activation in NF90-R cells. YAP signaling is known to be affected by cell density, therefore we examined YAP signaling at 10%, 50%, and 100% confluence. Sparse cell density typically induces YAP signaling by reducing pYAP levels, which results in YAP nuclear accumulation and increased expression of the YAP/TEAD target genes CTGF and Cyr61, while higher confluency results in cytoplasmic pYAP accumulation and CTGF/Cyr61 downregulation. As expected, expression of CTGF and Cyr61 was density dependent in the NF90 parental and S462 cell lines, however, pYAP expression was lost in the NF90-R cells, resulting in aberrant YAP activation and subsequent expression of CTFG and Cyr61 target genes, regardless of cell density (Figure 4C).

To investigate the impact of YAP/TEAD pathway inhibition on the resistant and sensitive lines, cells were plated at low and high densities and treated with a low or high dose of single agent pan-TEAD inhibitor, GNE-7883 (27). TEADi treatment strongly reduced CTGF expression in NF90-R cells, regardless of plating density (Figure 4D). There was no effect of GNE-7883 on ERK activation, and changes in AKT activation were variable and inconsistent across experimental replicates (Figure 4D). Collectively, these data suggest that long term exposure to trametinib uncouples density dependent repression of YAP signaling and maintains TEAD-dependent expression of target genes in MPNST to facilitate escape from contact inhibition.

To evaluate the efficacy of combined GNE-7883 and trametinib *in vitro*, the parental and resistant NF90 cell lines were plated at sparse and dense seeding densities and treated in a dose response matrix. Excitingly, GNE-7883 strongly sensitized the NF90-R cells to trametinib treatment, even at low doses (Figure 5A). Moreover, combination GNE-7883 and trametinib treatment profoundly reduced viability compared to single agent treatment in both cell lines, regardless of plating density (Figure 5A). This resulted in high Bliss synergy scores across multiple dose combinations in all experimental conditions (Figure 5A). These results demonstrate that combined TEAD and MEK inhibition is not only effective in trametinib resistant MPNST cells, but trametinib naïve cells as well. Interestingly, when we evaluated cell signaling upon GNE-7883 and trametinib treatment, we found that single agent trametinib was sufficient to reduce CTGF expression, with the strongest downregulation in combination treated cells (Figure 5B). These data suggest that crosstalk between the RAS/ERK and YAP/TEAD pathways occurs in MPNST, which could be exploited therapeutically.

**Figure 5:**
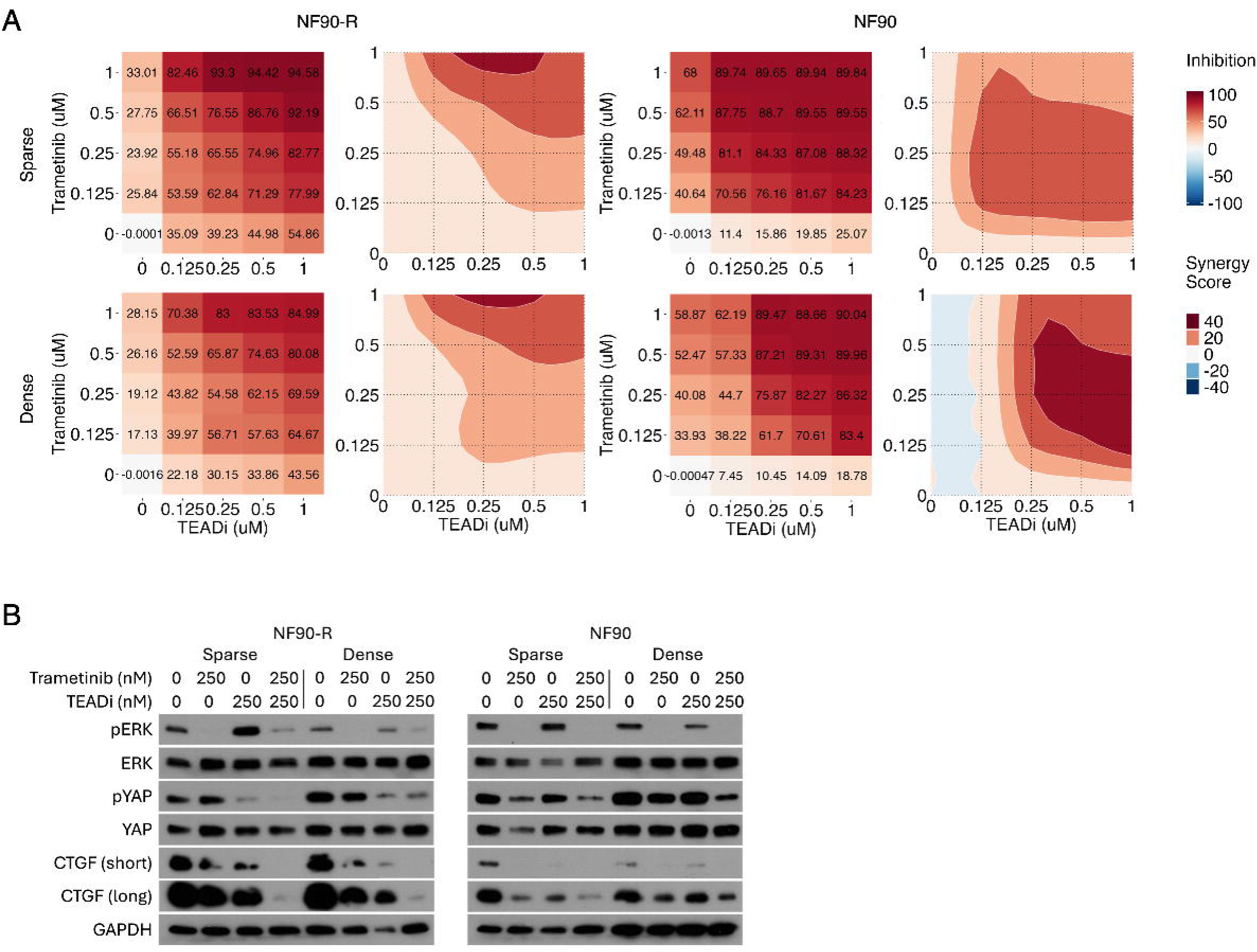
TEAD inhibition sensitizes MPNST cells to MEK inhibition in vitro A) Dose response matrices and corresponding synergy maps of NF90-R and NF90 percent inhibition and bliss synergy score in response to combined treatment with trametinib and TEADi at sparse (top) and dense (bottom) confluencies. B) Western blot of ERK and YAP/TEAD signaling in NF90-R and NF90 cells after combination TEADi and trametinib treatment for 24 hours.

### Transcriptional heterogeneity promotes resistance to combined TEAD and MEK inhibition *in vivo*

To test the efficacy of combination MEK and TEAD inhibition on tumor burden *in vivo*, we used the MEK inhibitors cobimetinib or trametinib plus the TEAD inhibitor GNE-7883 in the MPNST PDX tumor WU-356. There were no significant differences in tumor growth in single agent or combination treated animals compared to vehicle, although a subset of tumors did respond to combination treatment (Figure 6A), and tumor growth was significantly reduced in combination treated groups compared to single agent cobimetinib (Figure 6B). When we evaluated pathway inhibition in responder vs non-responder tumors, we found persistently high levels of both pERK and nuclear YAP in non-responders (Figure 6C, Supplemental Figure 5A), suggesting that pathway reactivation contributes to treatment resistance. We next hypothesized that nuclear YAP would be restricted to the edge of tumors to facilitate invasion, however IF revealed widespread nuclear YAP throughout tumors, even with TEADi treatment (Figure 6D, Supplemental Figure 5B), again indicating deregulation of YAP/Hippo dependent contact inhibition in MPNST.

**Figure 6:**
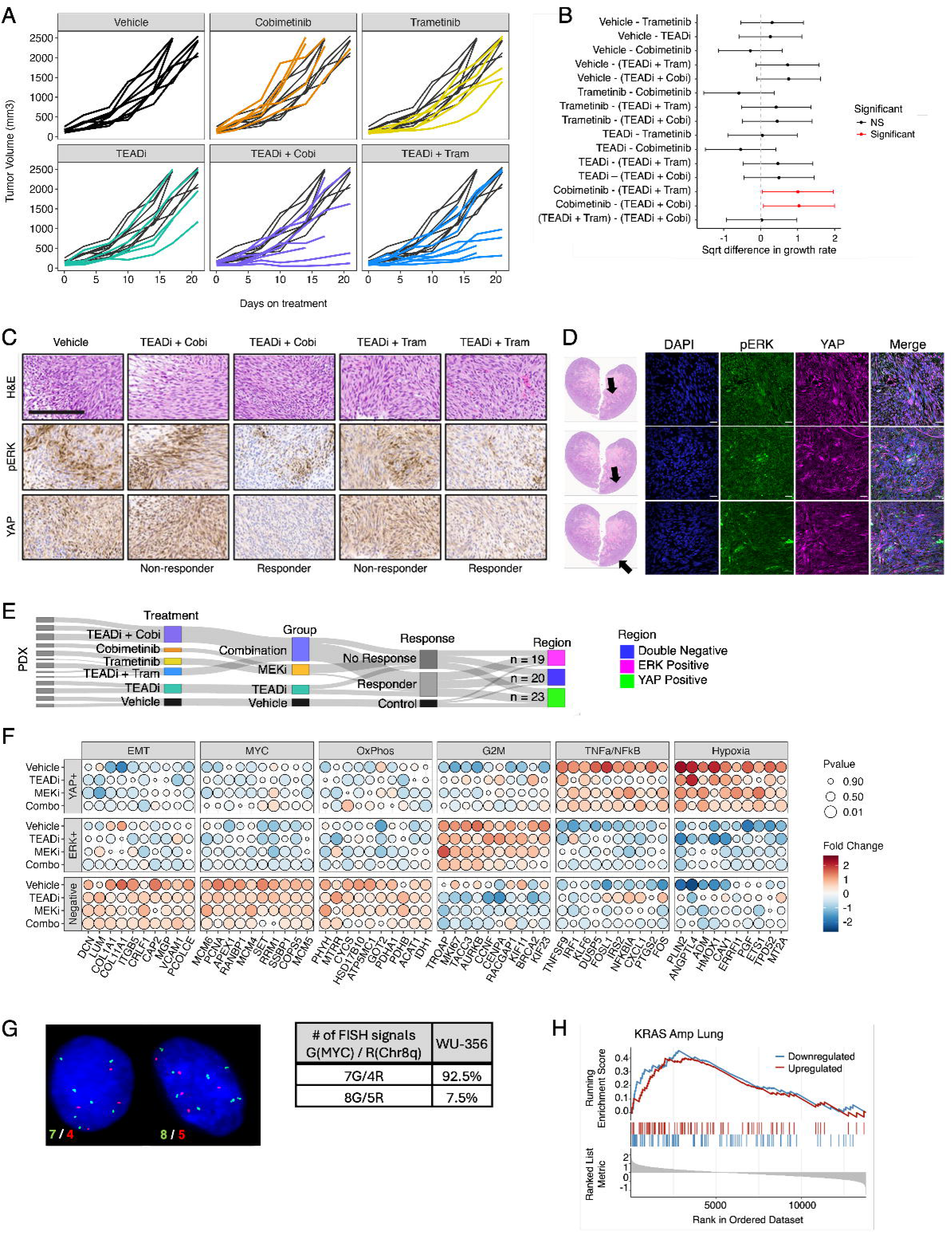
Combination MEKi and TEADi treatment response is heterogenous in vivo (A) Growth curves and B) statistical comparisons of WU-356 PDX tumorgrafts treated with GNE-7883 (TEADi), trametinib, cobimetinib, or a combination of TEADi and either trametinib or cobimetinib for 3 weeks or until tumors reach 2500 mm^3^. C) Representative immunostaining of pERK and YAP in combination treated tumorgrafts. D) Representative immunofluorescent staining of YAP in different tumor regions of TEADi treated tumor. E) Sankey diagram indicating the number of segmented regions sequenced, and the treatment groupings for downstream analysis. F) Dotplot of the top 10 core enrichment genes in each Hallmark pathway, stratified by segmented region and treatment group, sized by p value, and colored by log2 fold change in expression. G) Representative image and summary of FISH of MYC (green) and chromosome 8q (red) and signal ratios in the WU-356 PDX model. H) GSEA plot for KRAS gene sets in ERK positive regions compared to ERK negative regions.

We evaluated the transcriptional implications of ERK and YAP pathway reactivation using GeoMx, again selecting proliferative (Ki67 positive) regions for analysis. Due to tissue loss during processing, we combined cobimetinib and trametinib treated samples together as a single “MEKi” group for downstream analysis (Figure 6E). Contrary to our findings in the MSTA-440-2 PDX model in which we observed YAP-dependent EMT, EMT was enriched in the double negative population in WU-356 tumors (Figure 6F, Supplemental Table 2). Interestingly, this was in association with expression of MYC target genes (Figure 6F), suggesting that multiple transcriptional programs converge on EMT in MPNST, depending on the genomic context of the tumor. Indeed, MYC was highly amplified in the WU-356 PDX (Figure 6G). Several metabolic pathways, including oxidative phosphorylation, were also enriched in the double negative population (Figure 6F, Supplemental Table 2), while inflammatory and hypoxia pathways were enriched in YAP positive cells (Figure 6F, Supplemental Table 2), which was consistent with our findings in MSTA-440-2 PDXs (Figure 3H). Enrichment of G2M Checkpoint genes as well as deregulation of oncogenic signaling pathways predominated in the ERK positive population, with both positive and negative expression of RAS pathway genes (Figure 6F & H, Supplemental Table 2). This finding is intriguing, as we initially hypothesized that the RAS pathway would be exclusively turned on in the ERK positive population. We suspect that simultaneous up- and down-regulation of RAS effector transcriptional targets is due to rapid activation of both positive and negative feedback loops related to ERK/MAPK signaling in the absence of physiological RAS regulation by neurofibromin. Collectively these data indicate that inter- and intra-tumor heterogeneity contributes to treatment resistance and tumor progression/metastasis in MPNST.

## Discussion

*NF1-related* MPNSTs are highly resistant malignancies that are distinct from other soft tissue sarcomas due to their complex genomes and diminished survival (33). What is often underestimated about *NF1-related* MPNSTs is that they exhibit profound signaling plasticity that is not fully explained by putative genetic alterations. Up until recently, it has been difficult to quantify how intra- and inter-tumoral signal heterogeneity contributes to therapy resistance. Suitable analytical methods for dissecting tumoral heterogeneity are now beginning to emerge, such as scRNA seq (34, 35) and the spatial transcriptomics approach highlighted in this manuscript. These promising technologies should offer new insights into therapy resistance mechanisms and help identify promising therapeutic targets.

We now show that MPNST heterogeneity is not just a byproduct of genomic complexity. NF1-related MPNSTs exhibit kinome adaptation leading to the rapid development of resistant signaling clades even within genetically identical tumors. Our data confirm that therapy resistance mechanisms are not monolithic within a given tumor or even within a single PDX model (9). There exists a diverse number of signaling adaptations that drive therapy resistance to both single and combination kinase inhibitors. Interestingly, both inherent and adaptive resistance mechanisms were present in all our models, but there was substantial temporospatial and intra-tumoral heterogeneity among tumors with identical genetic backgrounds. These resistance mechanisms were both diverse and highly plastic, evolving within days in response to drug treatment or withdrawal (i.e. “drug holiday). Even when the *in vitro* data were promising (e.g. synergistic MEKi-TEADi response) therapy resistance was more commonly observed than therapy responses.

MPNSTs are defined by deregulation of the RAS pathway as a result of *NF1* deficiency, yet targeted treatments in both RAS and the parallel PI3K/AKT pathways have failed to demonstrate efficacy in MPNSTs (10, 36, 37). Here, we confirm that targeting and co-targeting AKT directly is ineffective in mitigating MEKi resistance despite strong evidence to suggest AKT inhibition should be effective (5). These negative data are important to communicate to the sarcoma research field but should not mitigate future attempts to target AKT directly as new drugs emerge. MEK inhibitor resistance remains, in part, AKT dependent, but targeting mTOR with everolimus showed better activity than directly targeting AKT. These results suggest that, although AKT activation is a common mechanism of MEKi resistance, there does not appear to be signaling dependency upstream of mTOR. To better understand how MPNST therapy resistance evolves, we induced signaling plasticity using a modified tumor kinetics approach. We leveraged genomically diverse PDX models using a drug holiday treatment scheme. Our intent was to induce kinome adaptation by short term therapy exposure and withdrawal using clinically available kinase inhibitors. While combined MEK and mTOR inhibition did initially show significant activity in a single PDX model, the drug holiday paradigm effectively induced resistance, as we did not see a statistically significant decrease in tumor size with any of our treatment combinations at the study endpoint. To pinpoint targetable pathways that mediate resistance we assessed spatial-temporal changes in gene expression using the GeoMx spatial transcriptomics platform. Interestingly, YAP target genes were strongly induced by the drug holiday, suggesting Hippo signaling as a node of resistance, which we confirmed using a MEKi resistant MPNST cell line.

In other RAS mutant or deregulated tumors, the Hippo pathway is now recognized as a key mechanism of resistance that offers an alternative target for treatment (20, 21, 32, 38). Furthermore, dysregulated YAP expression has been shown to cause malignant transformation of MPNST pre-cursor Schwann cells, in both a mouse and cell line model (17, 25). MPNSTs have also been identified as having the highest expression of active YAP across soft tissue sarcomas (16). Together these studies indicate that the Hippo signaling pathway could be providing a route of both tumor initiation and therapy resistance for MPNSTs, specifically through YAP activation. Here, we provide further evidence that MEK inhibitor resistance in MPNST is YAP/TEAD dependent. We also show for the first time, *in vitro* effectiveness of TEAD inhibition in MPNSTs. In our PDX model system, there was not a statistically significant response of combination MEK and TEAD inhibition among all grafted tumors. There was, however, a subset of that PDXs did respond well to combination treatment, suggesting the presence of distinct clonal populations present in the WU-356 PDX model, with varying dependency on MEK and YAP/TEAD pathways. We observed similar response heterogeneity in MSTA-440-2 PDXs that were treated with MEK and mTOR inhibitors. Collectively, these data imply that the genetic diversity of MPNSTs at diagnosis likely precludes treatment with single agent targeted therapies, including drugs that target the Hippo pathway. It also highlights the importance of studying TEAD inhibitors as a combination agent in the future.

Despite the lack of overall statistical significance, identifying RAS-Hippo-dependent “responders” is a critical step in targeting MPNSTs. We show that Hippo-dependent, MEKi-resistant signaling clades can be targeted despite profound background heterogeneity. It is interesting that statistical averaging would have otherwise masked these findings. For highly resistant tumors like MPNSTs, average tumor size may not always be a reliable way to define treatment effects when substantial heterogeneity is present. Examining kinase signaling differences between responder and non-responder tumors is a powerful way to identify new therapy targets and resistance mechanisms. The importance of Hippo signaling in MPNST progression has been previously reported by others (17, 25). Our work extends these important findings towards further testing of Hippo-targeted drugs in MPNSTs, such as TEAD inhibitors.

To understand intra-tumoral signaling heterogeneity in MPNSTs, we evaluated our treated PDX tumors using the GeoMx spatial transcriptomics platform. We profiled active ERK, YAP, and double negative populations in MSTA-440-2 and WU-356 PDX models. Interestingly, the MYC pathway was enriched in the double negative populations in both PDX models, suggesting cooperation of ERK, YAP, and MYC driven transcriptional programs in MPNST. Surprisingly, the pathways activated by these transcriptional populations were largely conserved across tumors, regardless of PDX model and treatment group. Given the well-established inter- and intra-tumor genomic heterogeneity in MPNST (7, 8), we had expected expression of unique gene sets across PDX models and treatment paradigms. Instead, our data confirm the presence of highly conserved populations with exclusive transcription profiles which we refer to as signaling clades, implying that they are capable of both convergent and parallel evolution (Figure 7A). To explain this intriguing finding, we propose two possible models for the evolution of specific transcriptional populations in MPNST. In a convergent evolution model, ERK, YAP, and MYC driven transcriptional programs evolve independently across patients due to similar environmental pressures in the MPNST microenvironment (i.e. transcriptional populations arise during tumor progression). Alternatively, in a parallel evolution model, ERK, YAP, and MYC driven transcriptional populations evolve similarly across patients as they share a common cell of origin whose daughter cells cooperate to support a favorable tumor microenvironment (i.e. transcriptional populations are essential for tumor initiation and are maintained throughout progression). Regardless of evolutionary mechanism, these distinct transcriptional populations direct growth, EMT, inflammation, and metabolic pathways in MPNST. Notably, expression of EMT genes was driven by YAP in the MSTA-440-2 PDX, but MYC in the WU-356 model, which was highly MYC amplified, suggesting that genomic alterations play a key role in transcriptional evolution in MPNST.

**Figure 7:**
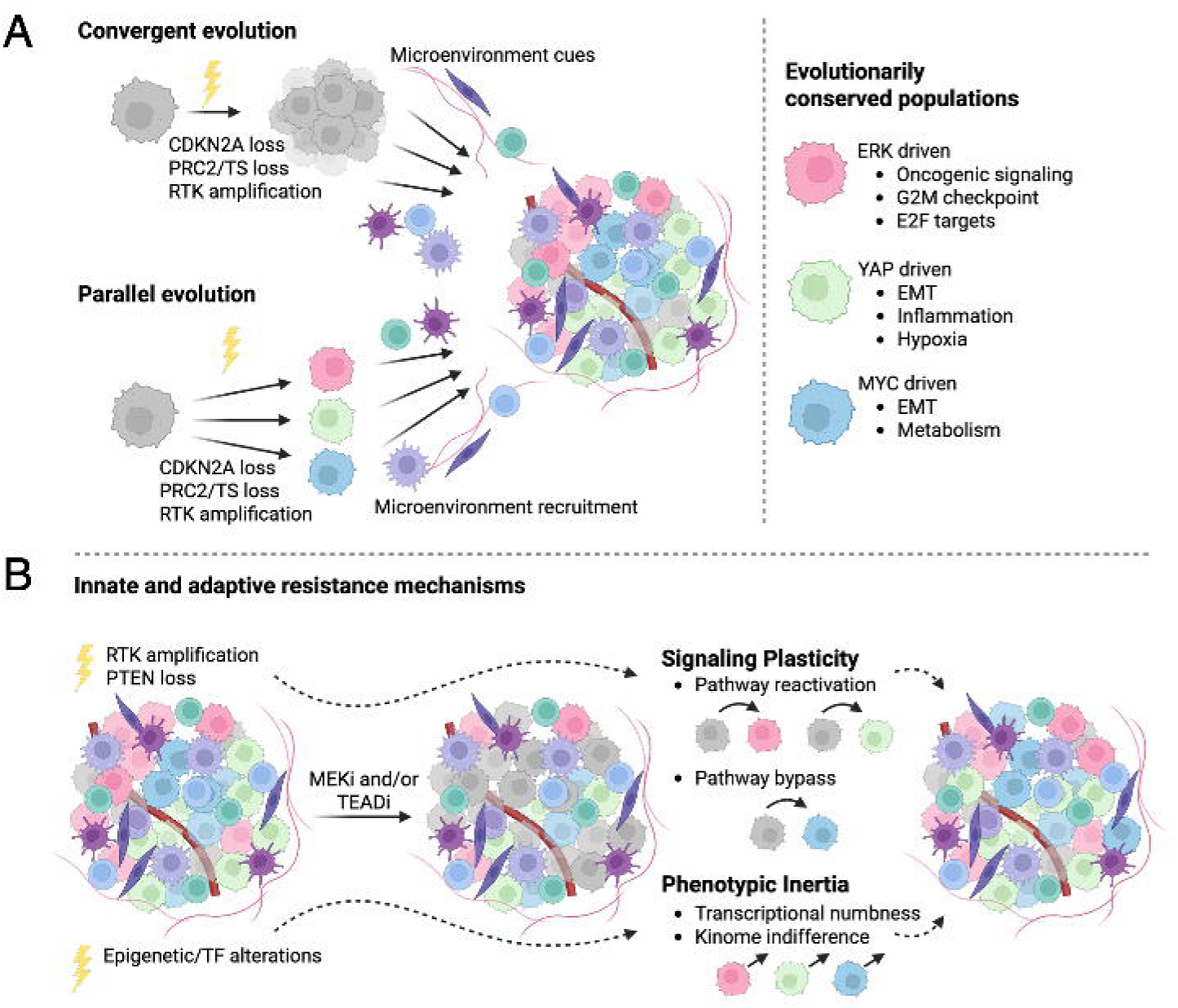
Proposed model of transcriptional evolution and subsequent drug resistance mechanisms A) Diverse genomic alterations result in conserved ERK, YAP and MYC transcriptional populations either via a convergent evolution model in which the post-initiation microenvironment triggers the development of these transcriptional populations as the tumor progresses, or via a parallel evolution model in which early intrinsic adaptation of these transcriptional programs directs tumor initiation. B) The diverse genomic alterations underlying conserved ERK, YAP, and MYC transcriptional populations contribute to distinct and redundant mechanisms of drug resistance. Genomic alterations in cell signaling regulators facilitate signaling plasticity via pathway reactivation or pathway bypass to reinstate the ERK, YAP, or MYC transcriptional programs upon MEK or TEAD inhibition. Meanwhile, genomic alterations in epigenetic regulators or transcription factors facilitate transcriptional numbness and kinome indifference in response to treatment, resulting in phenotypic inertia. Abbreviations: PRC2 polycomb repressive complex 2; TS tumor suppressor; RTK receptor tyrosine kinase; EMT epithelial to mesenchymal transition; TF transcription factor.

Unexpectedly, drug treatment had little impact on the ERK, YAP, or MYC transcriptional programs, with similar gene expression profiles across vehicle and drug treated tumors, regardless of drug response. These data suggest that even upon inhibition, ERK, mTOR, and Hippo driven transcriptional programs are maintained in MPNST. We previously identified signaling plasticity as a resistance mechanism in MPNST, by which oncogenic MAPK signaling pathways are reactivated via upstream RTK activation, or via pathway bypass in which the tumor cell switches to an alternative oncogenic pathway (6). In this study we identify an additional resistance mechanism, phenotypic inertia, which is the ability of cancer cells to tolerate stressful environments by ignoring cellular stress cues to maintain gene expression, resulting in enhanced tumor cell fitness. This “transcriptional numbness” is caused by loss of the normal epigenetic regulators of gene expression and provides tumor cells with a competitive advantage at the population level (39). Phenotypic inertia is predicted to play a critical role in MPNST cell stemness and tumor evolution (40) due to frequent alterations in the epigenetic regulators SUZ12 and EED, which are components of the histone methyltransferase complex PRC2 (41, 42). Consistent with this, we previously showed that resistance to MEK and MET inhibition was associated with kinome indifference in a genetically engineered mouse model of MPNST, where several resistant tumors showed almost no change in kinase activation upon drug treatment (6). Taken together, these data point to comprehensive and overarching drug resistance mechanisms in MPNST, by which tumors have evolved to rapidly adapt to cellular stress (plasticity) and ignore cellular stress (phenotypic inertia) upon drug treatment to maintain growth and metastasis pathways. We propose that genomic alterations in oncogenic signaling regulators facilitate signaling plasticity, while alterations in epigenetic regulators and transcription factors direct phenotypic inertia. In addition, heavy and diverse intra-tumoral mutational burden of MPNST ensures therapy resistance via multiple mechanisms that converge on sustained MAPK, Hippo, and MYC directed transcriptional programs (Figure 7B).

In the last decade, MEK inhibitors have revolutionized treatment for NF1 related plexiform neurofibromas, although they are ineffective in preventing or treating MPNST. Understanding how MEK inhibition alters signaling and transcriptional programs in the context of NF1 loss is critical for the development of new treatments for NF1-related tumors. Here we show that the mTOR and YAP/TEAD pathways are activated by MEK inhibitor exposure, yet blockade of these pathways in resistant MPNST PDX models does not significantly reduce tumor growth, despite strong *in vitro* synergy between trametinib and the novel TEAD inhibitor, GNE-7883. Using spatial transcriptomics, we uncovered phenotypic inertia as a key mechanism of drug resistance in MPNST, in addition to signaling plasticity. Further, we found that resistance is mediated by sustained ERK, YAP, and MYC driven transcriptional programs. In the future, preclinical studies should focus on addressing intratumoral heterogeneity and how it evolves over time.

## Methods

### Sex as a biological variable

Because of housing constraints, all mice used in this study were female. The findings of this study are not expected to be influenced by sex.

### Cell culture and drugs

Human MPNST cell lines S462 and NF90 were maintained at 37 °C in 5% CO2. NF90 cells were cultured in Dulbecco’s Modified Eagle’s Medium (DMEM, Thermo Fisher Scientific) with 10% fetal bovine serum (FBS, Corning) and 1% Penicilin/Streptomycin (Pen/Strep, Thermo Fisher Scientific. S462 cell lines were cultured in Roswell Park Memorial Institute 1640 (RPMI, Thermo Fisher Scientific) with 10% FBS and 1% Pen/Strep. To generate NF90-R cells, NF90 cells were exposed to escalating doses of trametinib over 4 months, starting at 10 nM, until the IC50 reached 800 nM (compared to 100 nM for the parental NF90 line). At this point, NF90-R cells were cultured with 2 nM of trametinib to retain selective pressure. All cell lines were tested for *Mycoplasma* infection every 6 months by PCR (ATCC). Trametinib (Novartis), afuresertib (Selleckchem), ipatasertib (Selleckchem), everolimus (Selleckchem), and GNE-7883 (Genentech) were prepared in DMSO, aliquoted, and stored at -80°C.

### Western blot

Cells were plated in 10 cm dishes at the specified cell densities, allowed to attach and grow for 24 hours, and then serum starved for 24 hours. Following serum starvation, cells were treated with the specified inhibitors for the described time. Cell lysates were collected in a RIPA buffer containing a protease/phosphatase inhibitor cocktail (Roche) and processed as previously described (11). Blots were incubated overnight at 4°C with the following primary antibodies from Cell Signaling Technology: pERK T202/Y204 (#9101, 1:5000), ERK (#9102, 1:5000), pAKT S473 (#9271, 1:1000), AKT (#9272, 1:2000), YAP (#14074, 1:1000), pYAP (13008, 1:1000), CTGF (#86651, 1:5000), CYR61 (#39382, 1:1000), and GAPDH (#2118, 1:5000).

### Viability assay

NF90 and NF90-R cell lines were plated at 5000 cells/well into a 96-well plate and allowed to adhere overnight. Cells were then treated with specified treatments at the indicated doses for 72 hours. Cell viability was measured using the CellTiter 96 Aqueous One Solution Cell Proliferation Assay (MTS, Promega) following the manufacturer’s protocol. Cell viability measurements were normalized to percent DMSO-treated control. Data were analyzed for Bliss synergy and plotted using the SynergyFinder package (v. 3.12.0) on R (v. 4.4.1) as described previously (43).

### Contact inhibition assay

NF90 and NF90-R cells were plated 5000 cells/well on top of Matrigel coating a 48 well plate. Cells were incubated at 37 °C in 5% CO2 and imaged every 24 hours for 72 hours.

### PDX animal studies

PDX MSTA-440-2 was derived from a patient at Helen Devos Children’s Hospital. PDX WU-356 were provided by Angela Hirbe at Washington University in St. Louis and previously characterized (8). Tumors were implanted into the flank of 6–8-week-old female NSG-SCID mice. For the drug holiday studies (Figure 2 and Supplemental Figure 2), once tumors reached approximately 150 mm^3^, the mice were randomized and placed on a treatment of 1 mg/kg trametinib, 5 mg/kg everolimus, or a combination of the two. Trametinib was suspended in a solution of 0.5% hydroxypropyl methylcellulose and 0.1% Tween 80. Everolimus was suspended in DMSO and PBS. Both compounds were administered via oral gavage 5 times weekly. Tumors were monitored twice a week and measured with calipers. After receiving treatment for 3 weeks, mice were placed on a “drug holiday” without treatment. Once drug holiday tumors reached 1100-1500 mm^3^, a 3 mm biopsy was taken of the tumor and mice were again randomized onto a second treatment of either trametinib, everolimus, or a combination of the two. Animals were euthanized when tumors reached 2500 mm^3^ or had received 15 days of treatment and tumors harvested for further analysis.

For the MEKi/TEADi study (Figure 6), WU-346 PDX tumors were implanted into the flank of 6–8-week-old female NSG-SCID mice. Once tumors reached approximately 150 mm^3^, the mice were randomized and placed on a treatment of 1 mg/kg trametinib, 5 mg/kg cobimetinib, 250 mg/kg GNE-7883 or a combination of the trametinib + GNE-7883 or cobimetinib + GNE-7883. Trametinib and cobimetinib were suspended in a solution of 0.5% hydroxypropyl methylcellulose and 0.1% Tween 80 administered via oral gavage Monday, Wednesday, and Friday. GNE-7883 was suspended in 100% sunflower seed oil and injected subcutaneously on an alternating schedule of 2 days on and 1 day off. Tumors were monitored twice a week and measured with calipers. After receiving treatment for 15 days, or once tumors reached 2500 mm^3^, mice were euthanized, and tumors harvested for further analysis.

To compare tumor growth, robust linear mixed-effects models were used to analyze the square-root transformed data. Data were analyzed in two groups according to drug delivery method. An interaction term for treatment and time was used as well as a random slope for time and a random intercept for each animal. The two vehicle groups (originally stratified by drug delivery method) were combined into a single group. R package emmeans was used to perform post-hoc comparisons. All p-values were false discovery rate corrected and an alpha of 0.05 was used. For the MEKi/mTORi study, treatment 1 data were loess smoothed and plotted using the stat_smooth function in the ggplot2 package with span set to 1. All analyses were conducted using R v4.1.0.

### Histology

Tumors were fixed in 10% neutral buffered formalin for 72 hours and then stored in 70% ethanol before undergoing routine histology processing and embedded in paraffin.

Deparaffinization, antigen retrieval, and staining were performed as previously described (44). Samples were stained with the following antibodies from Cell Signaling Technology: (IHC) pERK (#4370, 1:600), YAP (#14074, 1:100), and pS6 (#2215, 1:100); (IF) pERK (#5726, 1:200), and YAP-647 conjugated (#38707, 1:40).

### GeoMx

Tumor array tissue and slide processing, library generation, and sequencing for digital spatial profiling on the Nanostring GeoMx were performed by the Van Andel Institute Histology and Genomics Cores. A tissue microarray (TMA) was constructed using the formalin-fixed, paraffin-embedded tumors, with two areas per tumor represented in the TMA as 1 mm cores. The 5 um TMA sections were stained with the following antibodies: Ki-67-AF647 conjugated (CST #12075), YAP-AF488 conjugated (CST #14729, 1:100), and pERK (CST #4370, 1:100) (AF594 secondary), and labeled with UV photocleavable indexed oligo Human Whole Transcriptome Atlas probes and Syto83 nucleic acid (Thermo Fisher Scientific). The tissue areas of interest (AOIs) were located using fluorescent imaging on the GeoMx instrument. Regions of interest (ROIs) were selected and segmentation was performed based on antibody and DNA staining for desired tissue characteristics. Syto83 was used to ensure nuclei counts were high for collection. AOIs were processed sequentially on the instrument using UV light focused through each AOI releasing the photocleavable oligos into wells of a 96 well plate. After AOI collection, indexing primers were hybridized during Nanostring library preparation. Quality and quantity of the finished library pools were assessed using a combination of Agilent DNA High Sensitivity chip (Agilent Technologies, Inc.) and QuantiFluor® dsDNA System (Promega Corp.). Paired end, 50bp sequencing was performed on an Illumina Novaseq 6000 sequencer using an S2, 100 cycle sequencing kit v1.5 (Illumina Inc.). Basecalling was done by Illumina RTA3 and output was demultiplexed and converted to fastq format with bcl2fastq v1.9.0. Fastq files were then converted to Digital count conversion (DCC) files using GeoMx NGS Pipeline v2.3.3.10. Raw probe counts from the Human Whole Transcriptome Atlas panel underwent sequencing quality control (QC) using gene expression counts from each region of interest where regions that were found to be under-sequenced were dropped from further analysis. Probe QC identified each mRNA that was targeted by multiple probes and any outlier probes were subsequently removed from downstream analysis. The gene detection rate was greater than 15% in 184 of 190 ROIs and across all segment types, with 50% of segments having more than 7,000 detected genes. The remaining data underwent signal based Q3 normalization in NanoString GeoMx software, and individual counts were normalized against the 75th percentile of signal from their own areas of interest. Sankey diagrams were generated with the networkD3 (v0.4) R (v. 4.4.1) package. For differential expression contrasts, normalized data were fit to a linear mixed effect model using GeomxTools (v3.8.0). For the MSTA-440-2 drug holiday study, T3 vs T2 contrasts were initially generated for each treatment 2, with random intercepts to account for treatment 1, the animal ID, drug response, and segmented region, and all p values were adjusted using the BH method. GSEA was done using the clusterProfiler (v4.12.6) package (45). There was a strong overlap in pathway enrichment regardless of treatment 2 (Supplemental Figure 4), thus a summarized T3 vs T2 contrast was generated with treatment 2 as an additional random intercept to simplify the data visualization in Figure 3. Segment specific contrasts were also generated by treatment 2 for T3 MSTA-440-2 tumors, again controlling for animal ID and response. For the TEADi/MEKi treatment study in the WU-356 PDX, due to tissue loss during processing we first compared trametinib vs cobimetinib and (TEADi + trametinib) vs (TEADi + cobimetinib) groups. After p value adjustment, there were no significant differences in gene expression, thus, we combined trametinib and cobimetinib into a single MEKi group, and (TEADi + trametinib) and (TEADi + cobimetinib) into a single (TEADi + MEKi) group for downstream analysis. Segment specific contrasts for WU-365 tumors were generated by treatment group, with animal ID and response as random intercepts. Complete GSEA outputs are in Supplemental Tables 1 (MSTA-440-2) and 2 (WU-356). Dotplots were made using the ggpubr (v0.6.0) and ggplot2 (v3.5.1) packages. All analysis were done using R (v. 4.4.1).

### FISH

FISH probes were prepared from purified human BAC clones CTD-3056O22 (locus 8q24.21, MYC gene), CH17-311E13 and CH17-425G9 (locus 8q11.21 control), (BACPAC Resource Center, bacpac.chori.org). Locus 8q11.21 control clones were labeled with Orange-dUTP and 8q24.21 MYC clone (Abbott Molecular Inc., Abbott Park, IL), by nick translation. Tumor touch preparations were prepared by imprinting thawed tumors onto positively-charged glass slides. The sample slides were fixed in methanol:acetic acid (3:1) for 30 min and air-dried. Slides were then aged in 2X saline/sodium citrate (SSC) at 60 °C for 30 min, digested with 0.005% pepsin at 37 °C for 8 min, and washed with 1X PBS for 5 min. Slides were placed in 1% formaldehyde/PBS for 10 min at room temperature, washed with 1X PBS for 5 min, and dehydrated in an ethanol series (70%, 85%, 95%) for 2 min each. Slides were then denatured in 70% formamide/2X SSC at 74 °C for 3.5 min, washed in a cold ethanol series (70%, 85%, 95%) for 2 min each, and air-dried. The FISH probes were denatured at 75 °C for 5 min and held at 37 °C for 10-30 min until 10 ul of probe was applied to each sample slide. Slides were coverslipped and hybridized overnight at 37 °C in the ThermoBrite hybridization system (Abbott Molecular Inc.). The posthybridization wash was with 2X SSC at 73 °C for 3 min followed by a brief water rinse. Slides were air-dried and then counterstained with VECTASHIELD mounting medium with 4’-6-diamidino-2-phenylindole (DAPI) (Vector Laboratories Inc., Burlingame, CA). Image acquisition was performed at 600x system magnification with a COOL-1300 SpectraCube camera (Applied Spectral Imaging-ASI, Vista, CA) mounted on an Olympus BX43 microscope. Images were analyzed using FISHView v7 software (ASI) and at least 200 interphase nuclei were scored for each sample.

### Statistics

To compare tumor growth, robust linear mixed-effects models were used to analyze the square-root transformed data. Data were analyzed in two groups according to drug delivery method. An interaction term for treatment and time was used as well as a random slope for time and a random intercept for each animal. The two vehicle groups (originally stratified by drug delivery method) were combined into a single group. R package emmeans was used to perform post-hoc comparisons. All p-values were false discovery rate corrected and an alpha of 0.05 was used. For the MEKi/mTORi study, treatment 1 data were loess smoothed and plotted using the stat_smooth function in the ggplot2 package with span set to 1. All analyses were conducted using R v4.1.0.

For differential expression contrasts, normalized data were fit to a linear mixed effect model using GeomxTools (v3.8.0). For the MSTA-440-2 drug holiday study, T3 vs T2 contrasts were initially generated for each treatment 2, with random intercepts to account for treatment 1, the animal ID, drug response, and segmented region, and all p values were adjusted using the BH method. GSEA was done using the clusterProfiler (v4.12.6) package (45). There was a strong overlap in pathway enrichment regardless of treatment 2 (Supplemental Figure 4), thus a summarized T3 vs T2 contrast was generated with treatment 2 as an additional random intercept to simplify the data visualization in Figure 3. Segment specific contrasts were also generated by treatment 2 for T3 MSTA-440-2 tumors, again controlling for animal ID and response. For the TEADi/MEKi treatment study in the WU-356 PDX, due to tissue loss during processing we first compared trametinib vs cobimetinib and (TEADi + trametinib) vs (TEADi + cobimetinib) groups. After p value adjustment, there were no significant differences in gene expression, thus, we combined trametinib and cobimetinib into a single MEKi group, and (TEADi + trametinib) and (TEADi + cobimetinib) into a single (TEADi + MEKi) group for downstream analysis. Segment specific contrasts for WU-365 tumors were generated by treatment group, with animal ID and response as random intercepts. Complete GSEA outputs are in Supplemental Tables 1 (MSTA-440-2) and 2 (WU-356). Dotplots were made using the ggpubr (v0.6.0) and ggplot2 (v3.5.1) packages. All analysis were done using R (v. 4.4.1).

### Study approval

Research subjects were enrolled for tumor tissue harvesting in accordance with standardized institutional review board protocols and practice, under an approved Corewell Health/Van Andel Research Institute IRB protocol (SH/VAI IRB#2014–295). Subjects provided written, informed consent to enroll in the study. All animal experimentation in this study was approved by the Van Andel Institute’s Internal Animal Care and Use Committee (XPA-19-04-001). Sample sizes were selected based on our previously published study of murine MPNST tumorgrafts (11).

## Data availability

The spatial transcriptomics data generated in this study will be deposited in NCBI’s Gene Expression Omnibus upon publication of this work and will be accessible through a GEO Series accession number.

## Author contributions

LEM and JLG contributed equally – LEM is listed first because of their key role in project conceptualization.

Conceptualization: LEM, JLG; CRG, MRG; Data Curation: JLG, IB, ZF; Formal Analysis: LEM, JLG, IB, EW, ZF; Funding Acquisition: LEM, JLG, CRG, MRS; Methodology: LEM, JLG, CJE, SA, EAT, LT, KB, JEK, JAK, MA, IB, EW, ZF; Project Administration: CRG; Resources: JZ, AD, ACH; Supervision: LEM, JLG, CRG, MRS; Visualization: LEM, JLG, EAT, EW; Writing – Original Draft Preparation: LEM, JLG, LT, KB, MA, IB, EW, ZF, CRG, MRS; Writing – Review and Editing: LEM, JLG, JEK, JAK, ACH, CRG, MRS

## Competing Interests

AD and JZ: Employed by Genentch Inc. and held stock in Roche Inc. at the time of their contributions to this work; ACH: Advisory board SpringWorks Therapeutics, Inc. And Aadi Biosciences; grant funding from Tango Therapeutics; license agreement with Boehringer-Ingelheim.

## Acknowledgements

The authors thank the VARI Vivarium (RRID:SCR_023211), Bioinformatics and Biostatistics (RRID:SCR_024762), Genomics (RRID:SCR_022913), and Pathology and Biorepository (RRID:SCR_022912) Cores, as well as Zach Madaj for advice on statistical analysis.

## Funding

Department of Defense NFRP (W81XWH-21-1-0224) (MRS), Children’s Tumor Foundation Young Investigator Award (CTF-2018-01-009) (JLG), NF Michigan (MRS)

## Supplemental Figure Legends

Supplemental Figure 1: mTOR but not AKT inhibition is synergistic with trametinib A) Synergy maps showing Bliss synergy scores for afuresertib and trametinib treated NF90 (left) and S462 (right) cell lines. B) Dose response matrices and C) synergy maps of ipatasertib and trametinib treated cells. D) Synergy maps showing Bliss synergy scores for everolimus and trametinib treated cells.

Supplemental Figure 2: Trametinib and everolimus drug holiday study in WU-356 PDX A) Individual growth curves of WU-356 PDX lines treated with 1 mg/kg trametinib, 5 mg/kg everolimus, or a combination. Each treatment is represented by the corresponding color (as represented in legend) and plotted against all the individual tumor growth curves (gray). B) Representative IHC of WU-356 PDX for pERK, YAP, and pS6 at T2 and T3 timepoints.

Supplemental Figure 3: Signaling heterogeneity in MSTA-440-2 PDX drug holiday model Additional representative IHC images of re-exposure and crossover treated tumors shown in Figure 2.

Supplemental Figure 4: GSEA of MTORi and MEKi treated MSTA-440-2 T3 tumors compared to T2 biopsies A) Barplots indicating NES and qvalue for select MSigDB Hallmark and Oncogenic pathways, stratified by drug 2. B) Barplots for MSigDB Oncogenic pathway RAS related signatures in which both upregulated and downregulated gene sets were enriched.

Supplemental Figure 5: ERK and YAP signaling in TEADi and MEKi treated WU-356 PDXs A) Representative IHC images of single agent treated tumors from Figure 6A. B) Additional representative IF images of regional YAP expression in TEADi and MEKi treated tumors from Figure 6A.

## Notes

### Summary of Updates

Revised figures and text for clarity.

