## Supplemental Figures for "YAP signaling promotes resistance to MEK and AKT inhibition in *NF1*-related MPNSTs"

### Slide 1
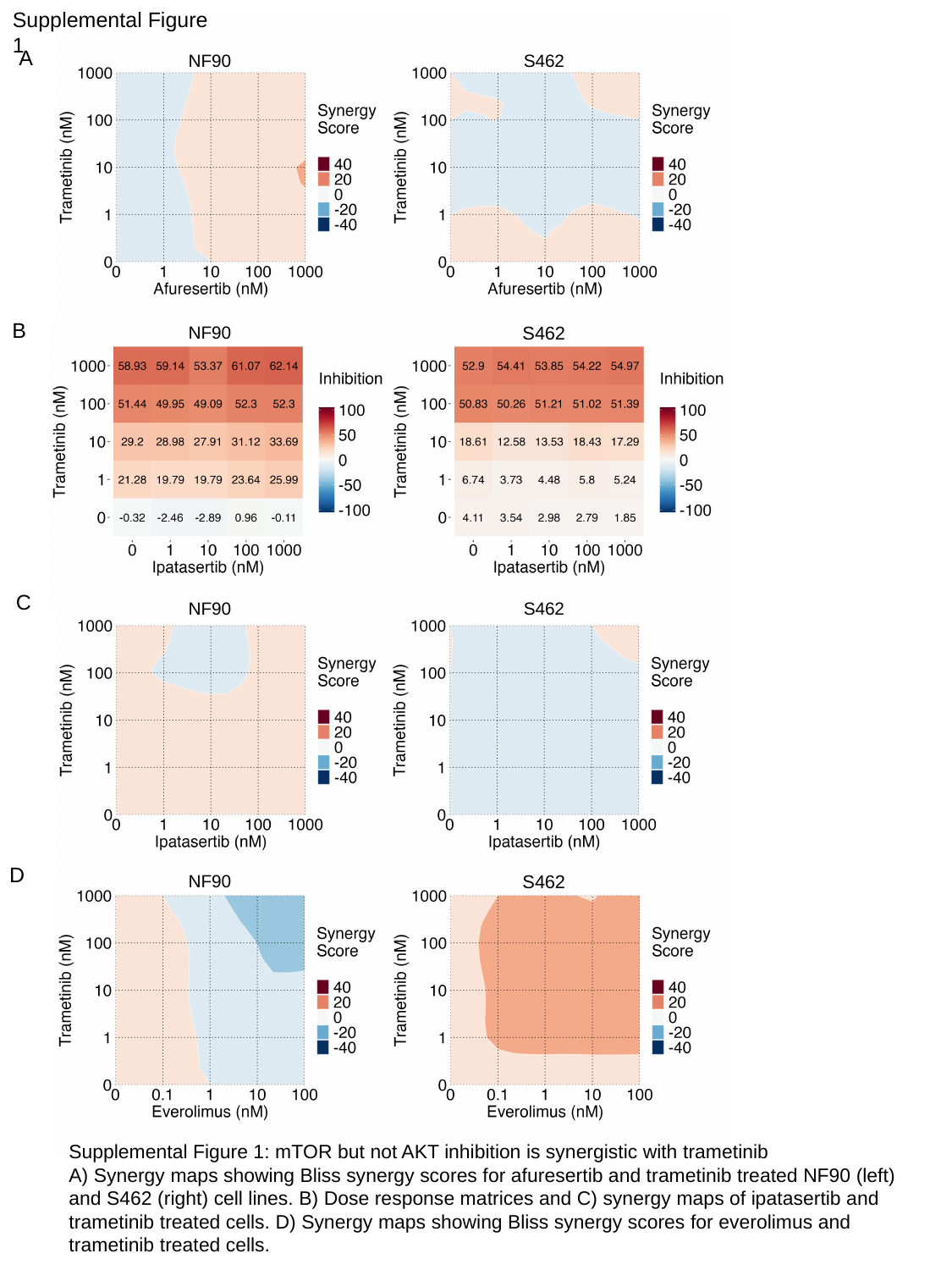

Supplemental Figure 1
A
NF90
S462
B
NF90
S462
C
NF90
S462
D
NF90
S462
Supplemental Figure 1: mTOR but not AKT inhibition is synergistic with trametinib
A) Synergy maps showing Bliss synergy scores for afuresertib and trametinib treated NF90 (left) and S462 (right) cell lines. B) Dose response matrices and C) synergy maps of ipatasertib and trametinib treated cells. D) Synergy maps showing Bliss synergy scores for everolimus and trametinib treated cells.

### Slide 2
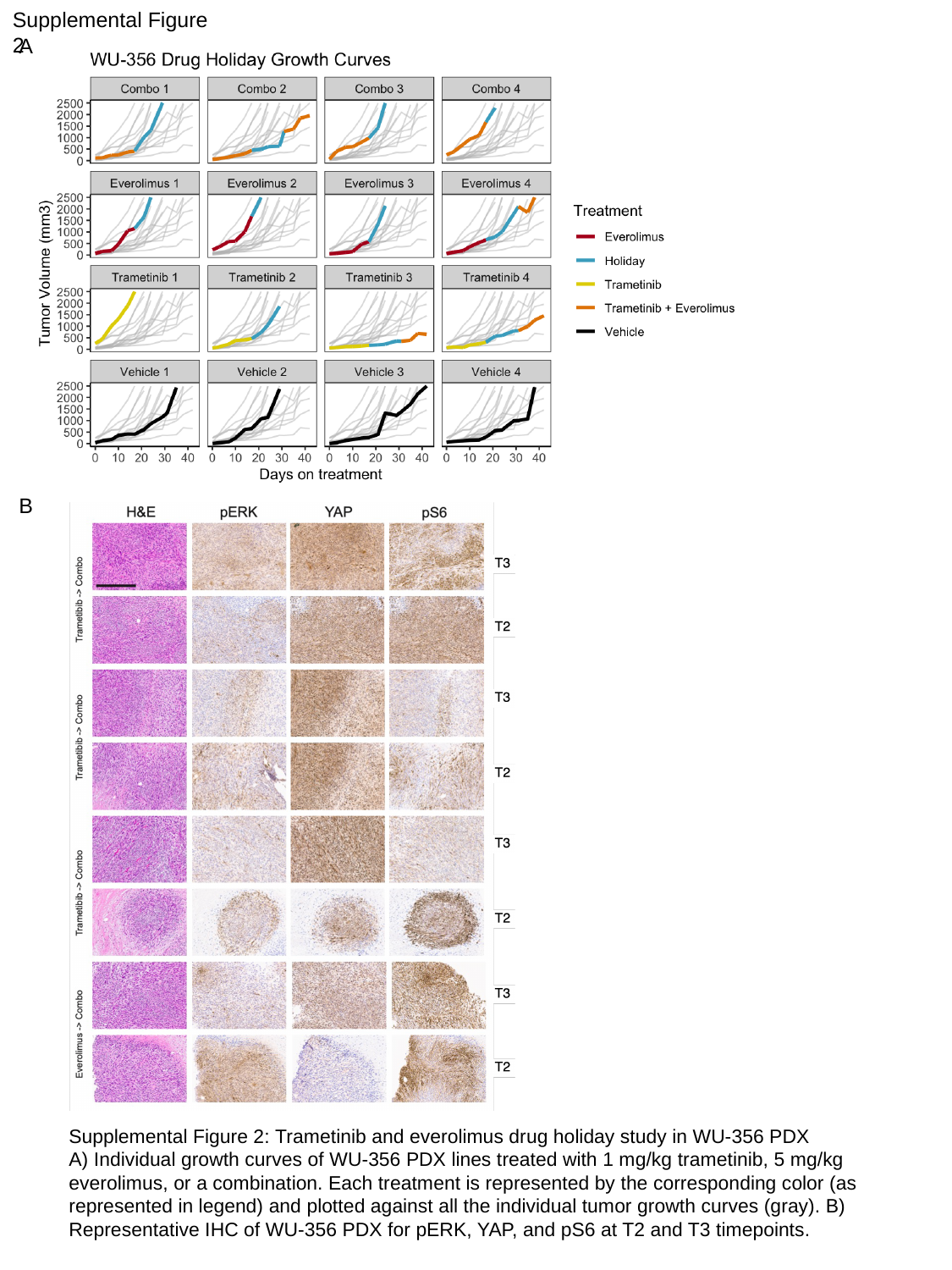

Supplemental Figure 2
A
B
Supplemental Figure 2: Trametinib and everolimus drug holiday study in WU-356 PDX
A) Individual growth curves of WU-356 PDX lines treated with 1 mg/kg trametinib, 5 mg/kg everolimus, or a combination. Each treatment is represented by the corresponding color (as represented in legend) and plotted against all the individual tumor growth curves (gray). B) Representative IHC of WU-356 PDX for pERK, YAP, and pS6 at T2 and T3 timepoints.

### Slide 3
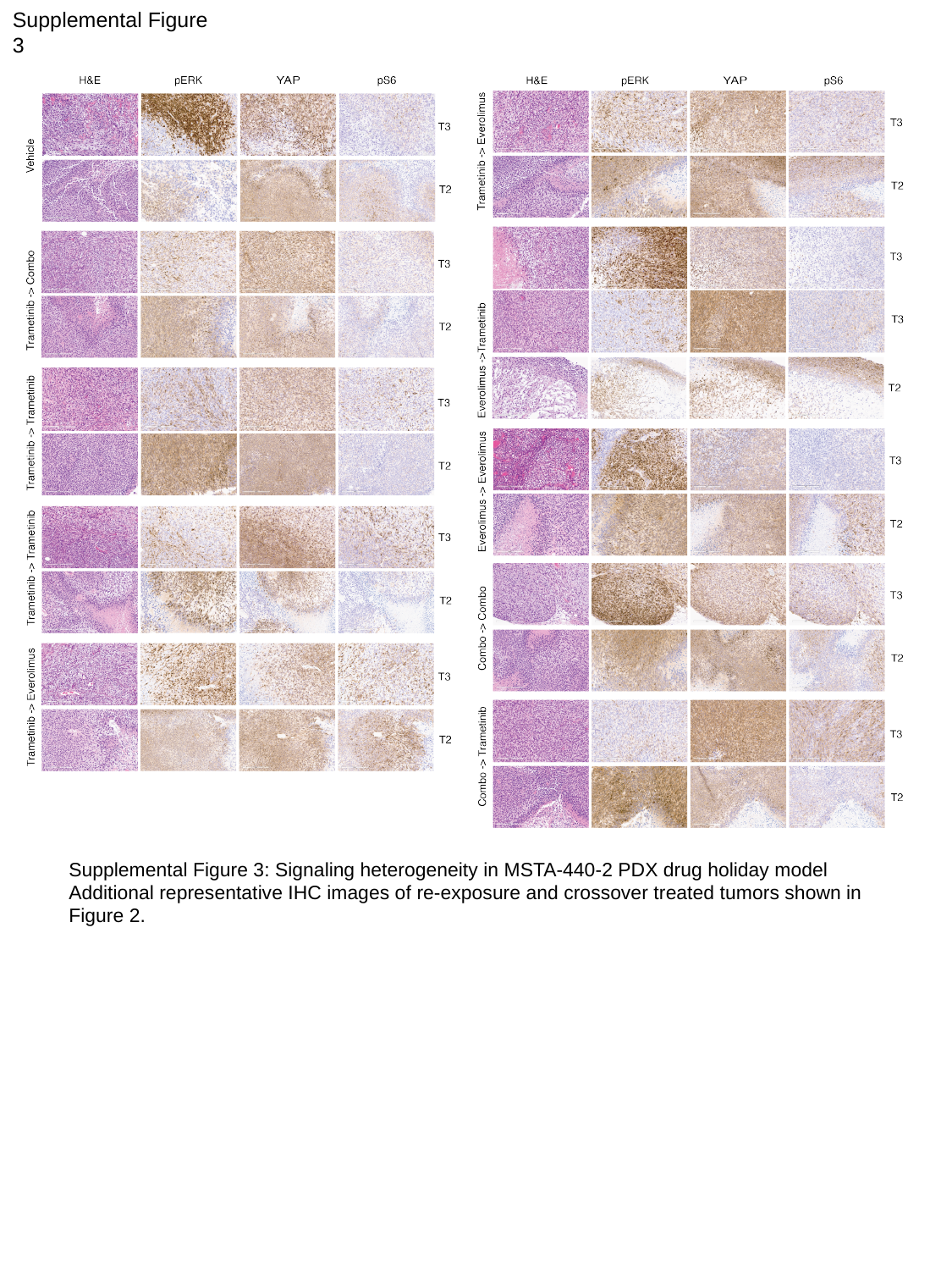

Supplemental Figure 3
Supplemental Figure 3: Signaling heterogeneity in MSTA-440-2 PDX drug holiday model
Additional representative IHC images of re-exposure and crossover treated tumors shown in Figure 2.

### Slide 4
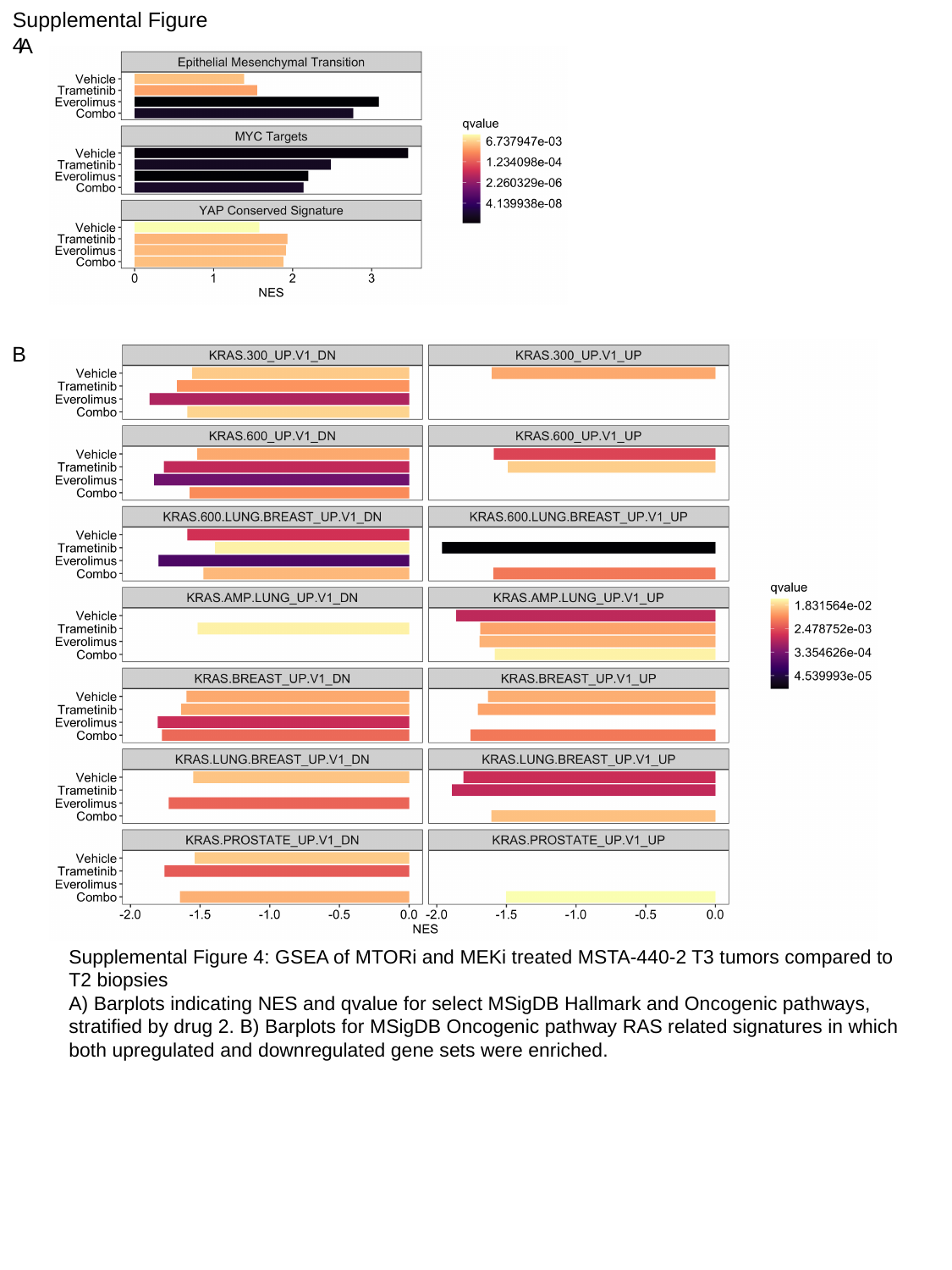

Supplemental Figure 4
A
B
Supplemental Figure 4: GSEA of MTORi and MEKi treated MSTA-440-2 T3 tumors compared to T2 biopsies
A) Barplots indicating NES and qvalue for select MSigDB Hallmark and Oncogenic pathways, stratified by drug 2. B) Barplots for MSigDB Oncogenic pathway RAS related signatures in which both upregulated and downregulated gene sets were enriched.

### Slide 5
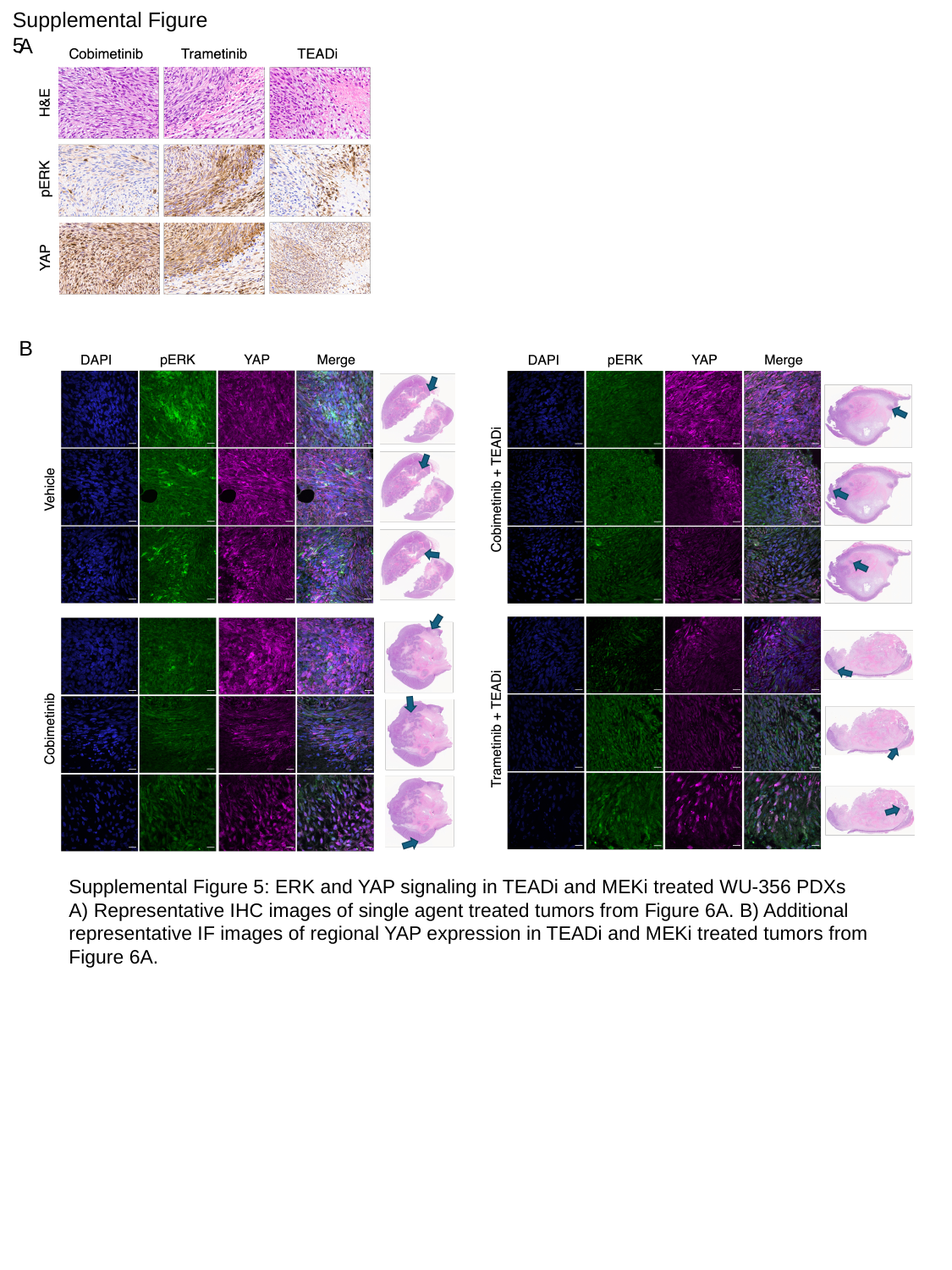

Supplemental Figure 5
A
B
Supplemental Figure 5: ERK and YAP signaling in TEADi and MEKi treated WU-356 PDXs
A) Representative IHC images of single agent treated tumors from Figure 6A. B) Additional representative IF images of regional YAP expression in TEADi and MEKi treated tumors from Figure 6A.
